## Supplementary figures and images for "An improved neural network model enables worm tracking in challenging conditions and increases signal-to-noise ratio in phenotypic screens"

### Figure S1

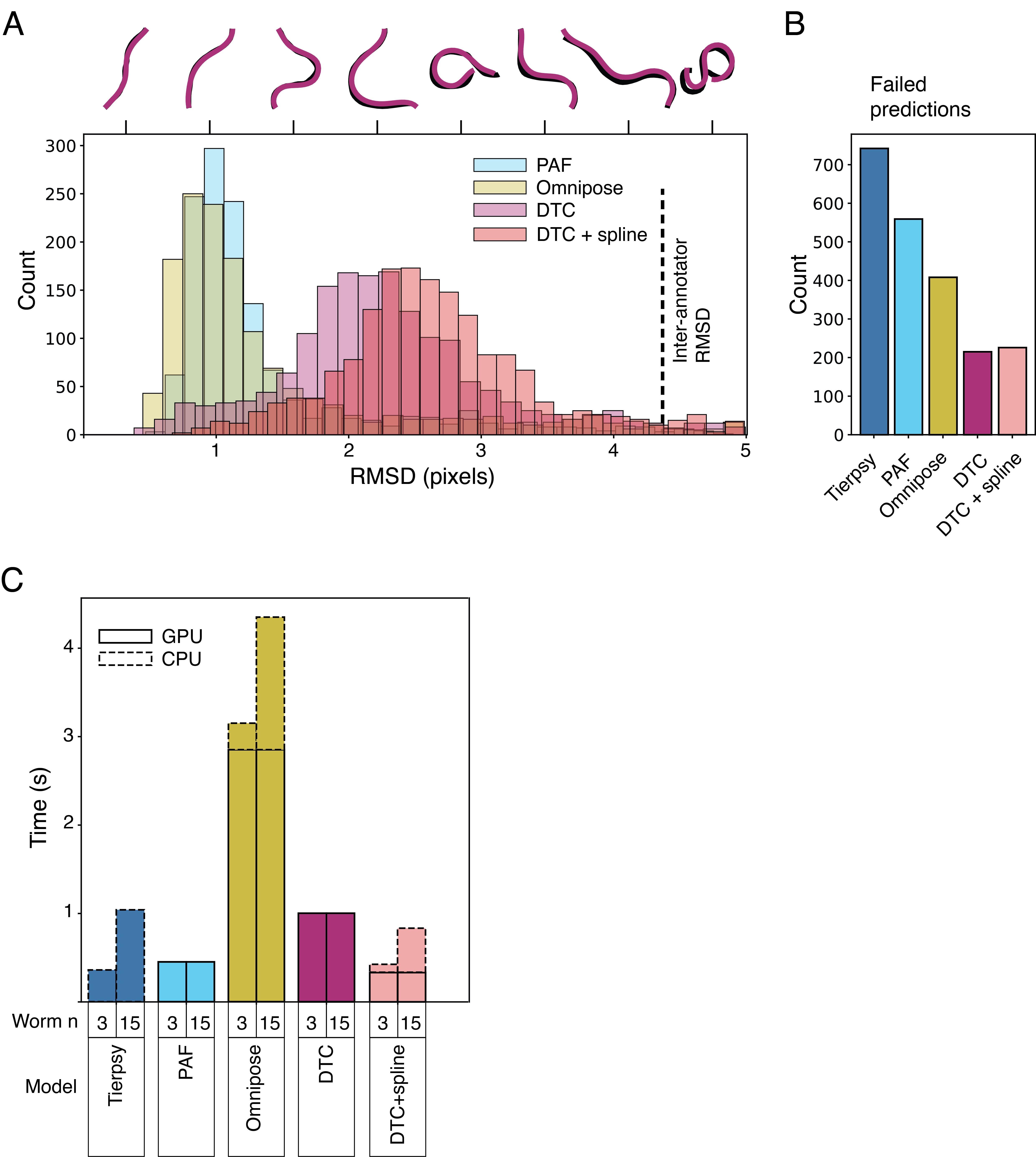
